# Epigenetic activation of meiotic recombination in Arabidopsis centromeres via loss of H3K9me2 and non-CG DNA methylation

**DOI:** 10.1101/160929

**Authors:** Charles J. Underwood, Kyuha Choi, Christophe Lambing, Xiaohui Zhao, Heïdi Serra, Filipe Borges, Joe Simorowski, Evan Ernst, Yannick Jacob, Ian R. Henderson, Robert A. Martienssen

**Affiliations:** Howard Hughes Medical Institute-Gordon and Betty Moore Foundation, Watson School of Biological Sciences, Cold Spring Harbor Laboratory, Cold Spring Harbor, NY, USA; Department of Plant Sciences, Downing Street, University of Cambridge, Cambridge, United Kingdom

**Keywords:** Meiosis, recombination, DSB, crossover, centromere, heterochromatin, DNA methylation, non-CG, H3K9me2, CMT3, Ds, Arabidopsis

## Abstract

Eukaryotic centromeres contain the kinetochore, which connects chromosomes to the spindle allowing segregation. During meiosis centromeres are suppressed for crossovers, as recombination in these regions can cause chromosome mis-segregation. Plant centromeres are surrounded by repetitive, transposon-dense heterochromatin that is epigenetically silenced by histone 3 lysine 9 dimethylation (H3K9me2), and DNA methylation in CG and non-CG sequence contexts. Here we show that disruption of Arabidopsis H3K9me2 and non-CG DNA methylation pathways increases meiotic DNA double strand breaks (DSBs) within centromeres, whereas crossovers increase within pericentromeric heterochromatin. Increased pericentromeric crossovers in H3K9me2/non-CG mutants occurs in both inbred and hybrid backgrounds, and involves the interfering crossover repair pathway. Epigenetic activation of recombination may also account for the curious tendency of maize transposon *Ds* to disrupt *CHROMOMETHYLASE3* when launched from proximal loci. Thus H3K9me2 and non-CG DNA methylation exert differential control of meiotic DSB and crossover formation in centromeric and pericentromeric heterochromatin.

## Introduction

Eukaryotic centromeres are the sites of kinetochore attachment to spindle microtubules that allow chromosome segregation (McKinley and Cheeseman 2015). Centromere identity is governed by nucleosomes containing CENPA/CENH3-related histone variants, which occupy large arrays of tandemly repeated satellite sequences (Malik and Henikoff 2009; Allshire and Karpen 2008). A conserved feature of centromeres shared across eukaryotes is suppression of crossover recombination during meiosis (Nambiar and Smith 2016; Lambie and Shirleen Roeder 1988; Copenhaver et al. 1999; Vincenten et al. 2015). Crossover suppression is important as centromere-proximal recombination events have been associated with chromosome segregation errors and aneuploidy (Koehler et al. 1996; Rockmill et al. 2006; Lamb et al. 1996). Plants, animals and fungi also typically possess repetitive pericentromeric heterochromatin, containing a high density of transposable elements (Malik and Henikoff 2009; Nambiar and Smith 2016; Allshire and Karpen 2008). In crop genomes, including wheat, barley and maize, these extensive pericentromeric heterochromatic regions occupy more than half of the chromosome and are also crossover-suppressed (Mayer et al. 2012; Choulet et al. 2014; Wei et al. 2009). However, the genetic and epigenetic factors that shape meiotic recombination patterns in eukaryote centromeric and pericentromeric regions remain to be fully understood.

Arabidopsis centromeres consist of megabase tandem arrays of the 178-180 base pair *CEN180* satellite repeat (Copenhaver et al. 1999; Kumekawa et al. 2000; Ito et al. 2007). The centromeric regions are densely DNA methylated and enriched in H3K9me2 and histone variant H2A.W (Stroud et al. 2013; Lippman et al. 2004; Lister et al. 2008; Yelagandula et al. 2014). Within the centromeric arrays, a subset of nucleosomes contain the centromeric histone H3 (CENH3) variant, which accumulate on specific *CEN180* variant sequences (Maheshwari et al. 2017; Nagaki et al. 2003). Surrounding the *CEN180* satellite arrays are repetitive, transposon-dense regions of pericentromeric heterochromatin (Stroud et al. 2013; Lippman et al. 2004; Lister et al. 2008; Yelagandula et al. 2014). Plant transposable elements are transcriptionally silenced by H3K9me2 and DNA cytosine methylation in CG and non-CG (CHG and CHH, where H=A, C or T) sequence contexts (Du et al. 2012; Stroud et al. 2013, 2014; Lippman et al. 2004). Arabidopsis mutants that lose maintenance of CG or non-CG DNA methylation have elevated transposon transcription and mobility at high and moderate levels, respectively (Stroud et al. 2014; Kato et al. 2003; Miura et al. 2001; Singer et al. 2001; Marí-Ordó ñ ez et al. 2013; Teixeira et al. 2009; Reinders et al. 2009; Tsukahara et al. 2012; Mirouze et al. 2009)

In Arabidopsis, the DNA chromomethyltransferases CHROMOMETHYLASE2 (CMT2) and CHROMOMETHYLASE3 (CMT3) recognize heterochromatic H3K9me2 via BAH and chromodomains, and methylate associated DNA in CHH and CHG contexts, respectively (Lindroth et al. 2001; Stroud et al. 2014, 2013; Du et al. 2012; Bartee et al. 2001; Zemach et al. 2013; Dubin et al. 2015). Methylation of histone H3K9 requires the SET domain methyltransferases KRYPTONITE/SUVH4, SUVH5 and SUVH6, which are recruited to methylated DNA by SRA methyl-cytosine binding domains (Du et al. 2014; Stroud et al. 2014; Johnson et al. 2007; Malagnac et al. 2002). In contrast, in fission yeast, which lacks DNA methylation, the SET domain histone lysine methyltransferase CRYPTIC LOCI REGULATOR4 (CLR4) is recruited to methylated H3K9me2 directly via its chromodomain (Allshire and Ekwall 2015). Thus, by separating chromodomains (CMT2 and CMT3) from SET domains (KYP/SUVH5/SUVH6), plants have introduced non-CG DNA methylation as an additional layer of epigenetic control underlying H3K9me2. The *de novo* DNA methyltransferase DOMAINS REARRANGED METHYLTRANSFERASE2 (DRM2) is also required for maintenance methylation of non-CG contexts, and thus can also impact upon H3K9me2 (Stroud et al. 2014, 2013; Cao and Jacobsen 2002; Cao et al. 2003). Alongside these mechanisms, the METHYLTRANSERFERASE1 (MET1) cytosine methyltransferase, VARIANT IN METHYLATION1 (VIM1) family proteins and the DECREASE IN DNA METHYLATION1 (DDM1) SWI/SNF chromatin remodeller are required for maintenance of CG context DNA methylation (Kankel et al. 2003; Saze et al. 2003; Lippman et al. 2004; Stroud et al. 2013; Lister et al. 2008; Woo et al. 2008). In this work we use mutations in Arabidopsis heterochromatic silencing pathways to investigate epigenetic control of meiotic recombination in the centromeric and pericentromeric regions.

Meiotic crossovers form via interhomolog repair of DNA double strand breaks (DSBs) that are generated by SPO11 topoisomerase-related complexes (Robert et al. 2016; Vrielynck et al. 2016; Keeney et al. 1997). Diverse eukaryotes show the presence of recombination hotspots, which are approximately kilobase regions with an elevated frequency of meiotic DBSs or crossovers, compared to the genome average or surrounding areas (Kauppi et al. 2004; Choi and Henderson 2015; de Massy 2013). Hotspots in different eukaryotic lineages are controlled to varying degrees by genetic and epigenetic information (Kauppi et al. 2004; Choi and Henderson 2015; de Massy 2013). At the chromosome-scale, Arabidopsis crossover frequency is highest in gene-dense euchromatin, whereas the heterochromatic centromeres are crossover-suppressed (Giraut et al. 2011; Salomé et al. 2012; Copenhaver et al. 1999; Yelina et al. 2015). At the fine-scale, plant crossover hotspots occur at gene promoters and terminators, and recombination is promoted by euchromatic modifications, including histone variant H2A.Z (Wijnker et al. 2013; Hellsten et al. 2013; Choi et al. 2013; Shilo et al. 2015). Acquisition of DNA methylation and H3K9me2 via the RNA directed DNA Methylation (RdDM) pathway is sufficient to silence Arabidopsis euchromatic crossover hotpots (Yelina et al. 2015). This is consistent with DNA methylation suppressing meiotic DNA double strand breaks in mouse (Zamudio et al. 2015), and silencing crossovers in Ascobolus (Maloisel and Rossignol 1998). Furthermore, loss of RNAi and CLR4-dependent H3K9me2 elevates centromeric crossovers in fission yeast (Ellermeier et al. 2010), and Drosophila position effect variegation (PEV) suppressor mutations (*Suppressor of Variegation*) can modify centromeric crossover frequency (Westphal and Reuter 2002). The Arabidopsis *met1* and *ddm1* mutants associate with remodelling of meiotic recombination along chromosomes, with crossover increases in the chromosome arms and decreases across the centromeres (Yelina et al. 2015, 2012; Melamed-Bessudo and Levy 2012; Colomé-Tatché et al. 2012; Mirouze et al. 2012). However, how non-CG DNA methylation and other epigenetic silencing pathways contribute to recombination landscapes along plant chromosomes has not been fully explored. In this study we address the roles of H3K9me2 and non-CG DNA methylation in suppression of meiotic DSBs and crossovers within Arabidopsis centromeres and pericentromeric heterochromatin.

## Results

### Transposon insertions into *CMT3* appear to induce meiotic recombination

In a large scale screen for *de novo* insertions of the non-autonomous maize transposable element *Dissociation* (*Ds*) introduced into Arabidopsis, a transgene strongly expressing the *Activator* (*Ac*) transposase was crossed to plants containing *Ds* ‘launch-pads’ triggering *Ds* transposition in clonal cell lineages within F_1_ plants (Sundaresan et al. 1995). As *Ds* elements are known to preferentially transpose to linked sites (Bancroft and Dean 1993; Jones et al. 1990), a positive-negative selection scheme was implemented in F_2_ progeny to select *against* the *Ds* launch-pad and *for* transposed *Ds* (Sundaresan et al. 1995). In this way recovery of F_2_ transpositions was dependent on recombination between the transposed *Ds* and the launch-pad, and should be mostly unlinked. Three launch-pads on chromosome 1, *DsE2, DsE3* and *DsE6* (Fig. 1), were used to generate 9,622 independent transpositions, which were mapped using TAIL-PCR (Springer et al. 1995). Unexpectedly, we recovered a dramatic enrichment of homozygous and heterozygous insertions into the *CMT3* locus, but only when *Ds* elements were launched from a closely linked (~ 360 kb) proximal locus on chromosome 1 (*DsE3*), and not from a distal locus (*DeE6*) located a comparable distance from *CMT3* (~ 624 kb) (Fig. 1). One possible explanation was that flowers carrying homozygous *cmt3* insertions arose by transposition and mitotic recombination, consistent with active *Ac* elements stimulating mitotic recombination at the maize *p1* locus (Athma and Peterson 1991; Xiao et al. 2000). Importantly, only mitotic recombination with proximal launch-pads, and not distal, would result in flowers homozygous for the transposed *Ds* element (Athma and Peterson 1991; Xiao et al. 2000). If *cmt3* causes increased meiotic recombination in vicinity to the *Ds* elements this would then promote separation from the launch-pad, and thereby lead to increased recovery of F_2_ transposant progeny carrying heterozygous or homozygous insertions that disrupt *CMT3*. Based on these observations we sought to further investigate the role of CMT3 and the non-CG/H3K9me2 pathway on Arabidopsis meiotic recombination.

**Figure 1.**
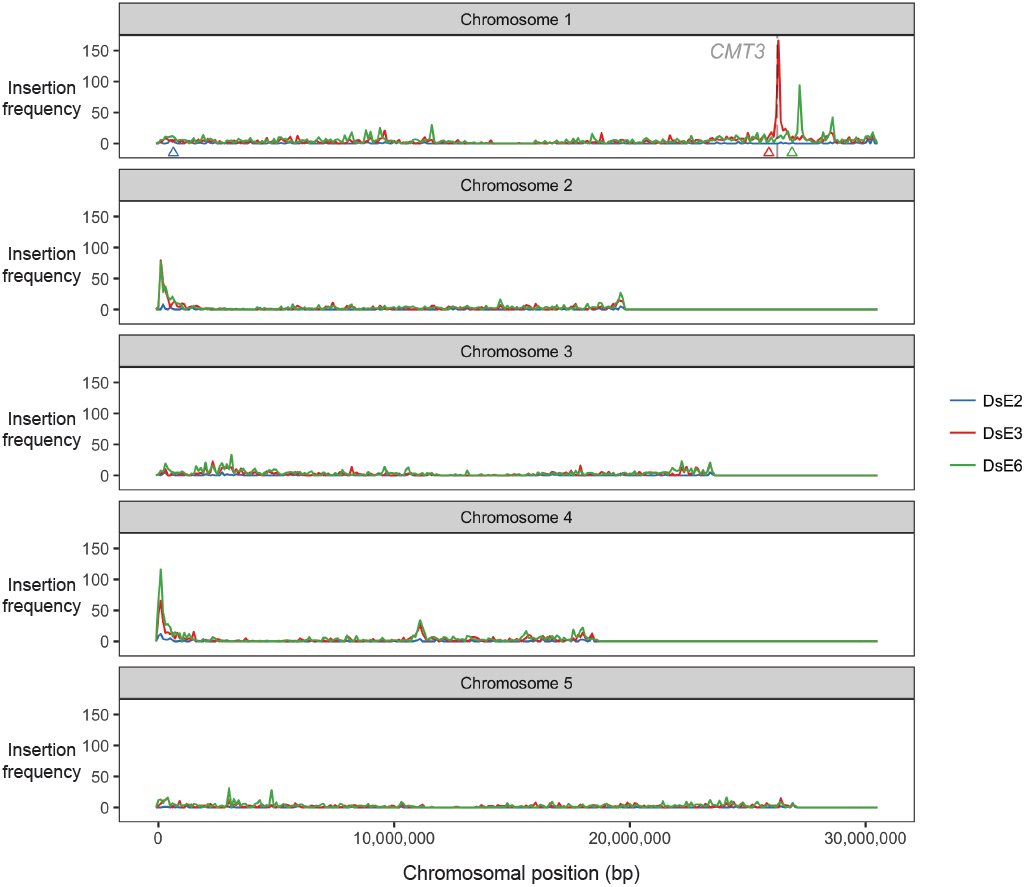
Transposon insertions in *CMT3* appear to induce recombination. Enhancer trap DNA transposons (*DsE*) based on the maize transposable element *Dissociation* (*Ds*) were introduced into the Arabidopsis genome on a T-DNA with a negative selectable marker (Sundaresan et al. 1995). Three independent *DsE* launch-pads were mapped to chromosome 1: *DsE2* (633,819 bp, blue triangle), *DsE3* (25,893,665 bp, red triangle) and *DsE6* (26,877,449 bp, green triangle). After introducing *Activator* transposase, 9,622 transpositions of *DsE* were isolated in F_2_ progeny by selecting for the transposon, but against the launch-pad, to select against linked transpositions which would otherwise be highly favored. New *DsE* insertions were mapped by sequencing flanking DNA. Transpositions from each chromosome 1 launch-pad are displayed in 100 kb bins, and most were unlinked, including two hotspots corresponding to nucleolar organizer regions at the ends of chromosomes 2 and 4. However, 2 much sharper insertions hotspots, on chromosome 1, were specific for closely linked launchpads, *DsE3* (red triangle) and *DsE6* (green triangle), respectively. The hotspot immediately distal to *DsE3* (~ 4% of transpositions launched from *DsE3*) was in and immediately around the *CMT3* gene (26,248,318 - 26,253,585 bp).

### Epigenetic activation of pericentromeric crossovers in non-CG/H3K9me2 mutants

To investigate meiotic recombination frequency in Arabidopsis non-CG DNA methylation and H3K9me2 pathway mutants, we used fluorescent crossover reporter lines (Fluorescent Tagged Lines, FTLs) (Berchowitz and Copenhaver 2008; Melamed-Bessudo et al. 2005). FTL lines express different colours of fluorescent protein under seed (*NapA*) or pollen (*LAT52*) specific promoters, from linked T-DNAs insertions (Fig. 2A). The scoring of fluorescent colour inheritance in the progeny seed or pollen (male gametes) of FTL hemizygotes allows the measurement of sex-averaged or male-specific crossover frequency, respectively, in defined chromosomal intervals (Fig. 2A) (Berchowitz and Copenhaver 2008; Melamed-Bessudo et al. 2005; Yelina et al. 2013).

**Figure 2.**
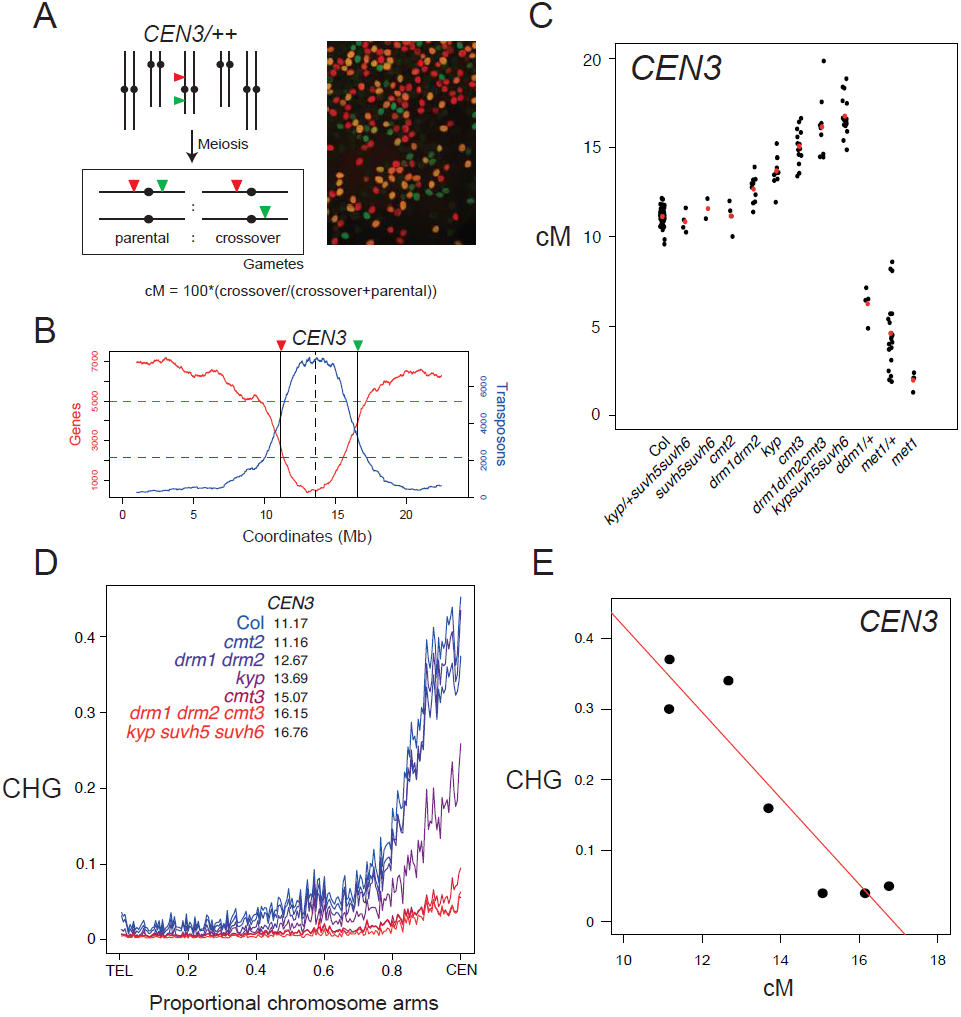
Progressive increases of pericentromeric crossover frequency in H3K9me2 and non-CG DNA methylation pathway mutants. **(A)** Measurement of crossover frequency using segregation of hemizygous Fluorescent Tagged Line (FTL) T-DNAs. A representative fluorescent micrograph is shown of *FTL/++* pollen, reproduced from (Choi et al. 2016). **(B)** Gene (red) and transposon (blue) density along chromosome 3, with the location of *CEN3* FTL T-DNAs indicated by vertical lines. Mean values are indicated by the horizontal dotted lines, and the centromere by the vertical dotted line. **(C)** *CEN3* crossover frequency (cM) in DNA methylation mutants. Data for *met1* and *met1/+* are reproduced from (Yelina et al. 2015). Black dots represent replicate measurements and red dots show mean values. **(D)** Published BS-seq data (Stroud et al. 2013) was used to analyse CHG DNA methylation density analysed along chromosome telomere-centromere axes in wild type and H3K9me2 and non-CG DNA methylation mutants. Lines are coloured according to *CEN3* cM (red=highest, blue=lowest). **(E)** Correlation between *CEN3* cM and CHG DNA methylation from published BS-seq data (Stroud et al. 2013).

We first analysed crossover frequency within the 5.4 megabase (Mb) *CEN3* FTL interval, which spans the centromere and pericentromeric heterochromatin of chromosome 3, in wild type versus non-CG/H3K9me2 mutants (Fig. 2B) (Yelina et al. 2015). Genetic ablation of the H3K9 methyltransferases (*kyp suvh5 suvh6*), or the non-CG DNA methyltransferases (*drm1 drm2 cmt2 cmt3*), eliminates both H3K9me2 and non-CG DNA methylation, while single and double mutants have intermediate effects (Stroud et al. 2014, 2013; Cao et al. 2003). We observed that mutations that disrupt H3K9me2 and non-CG DNA methylation to progressively greater extents resulted in progressively greater increases in *CEN3* crossover frequency (*suvh5 suvh6 < cmt2 < drm1 drm2 < kyp < cmt3 < drm1 drm2 cmt3 < kyp suvh5 suvh6*) (all *X*^*2*^ *P*< 2.0 × 10^−16^) (Fig. 2C and Supplemental Table S1) (Stroud et al. 2014, 2013).

We used published bisulfite sequencing data to analyse DNA methylation levels within the *CEN3* interval in the genotypes analysed for crossover frequency (Stroud et al. 2013). Within this series of mutants, levels of CHG DNA methylation showed a strong negative correlation with recombination (Pearson’s *r= −*0.93 *P* = 2.36 × 10^−3^), whereas CG and CHH methylation were not significantly correlated (Fig. 2D-2E and Supplemental Table S2) (Stroud et al. 2013). This is consistent with the CHG/H3K9me2 pathway inhibiting meiotic crossovers in the centromeric regions. In contrast to non-CG/H3K9me2 mutants, heterozygous *ddm1/+, met1/+* or homozygous *met1* mutants, which inherit chromosomes hypomethylated in the CG context, have reduced *CEN3* recombination, as reported previously (Fig. 2C and Supplemental Table S1) (Yelina et al. 2012, 2015; Mirouze et al. 2012; Melamed-Bessudo and Levy 2012; Colomé-Tatché et al. 2012). Thus, loss of the CG versus non-CG DNA methylation maintenance pathways causes distinct effects on pericentromeric crossover recombination, in addition to different effects on transposon transcription and mobility (Stroud et al. 2012; Yelina et al. 2015; Mirouze et al. 2012; Melamed-Bessudo and Levy 2012; Colomé-Tatché et al. 2012; Kato et al. 2003; Miura et al. 2001; Stroud et al. 2013, 2014; Zhang et al. 2006; Lippman et al. 2004; Lister et al. 2008; Henderson and Jacobsen 2008).

We confirmed release of crossover suppression in *cmt3* across the chromosome 5 centromere and pericentromeric heterochromatin, using additional FTLs in Col (*CTL5.11, X*^*2*^ *P=*1.30 × 10^−3^) and Ler (*LTL5.4, X*^*2*^ *P*=2.09 × 10^−9^) inbred backgrounds (Supplemental Fig. S1 and Supplemental Tables S3 and S4). We also crossed *cmt3* alleles in Col (*cmt3-11*) and Ler (*cmt3-7*) accessions together to generate Col/Ler F_1_ that were *cmt3* mutant and carried the *CEN3* FTL (Fig. 3A) (Lindroth et al. 2001). We observed significantly increased *CEN3* genetic distance in *cmt3* hybrids (*X*^*2*^ *P=*1.27 × 10^−86^), similar to the increase observed for inbreds (Fig. 3B and Supplemental Tables S1 and S5). In contrast, recombination in the euchromatic *420* FTL interval on chromosome 3 did not significantly change in *cmt3* inbreds, and slightly decreased in hybrids (*X*^*2*^ *P=*1.96 × 10^−3^), compared to wild type (Fig. 3C and Supplemental Table S6). Together these data indicate that mutations in the H3K9me2/non-CG pathway primarily activate crossover frequency in proximity to the centromeres.

**Figure 3.**
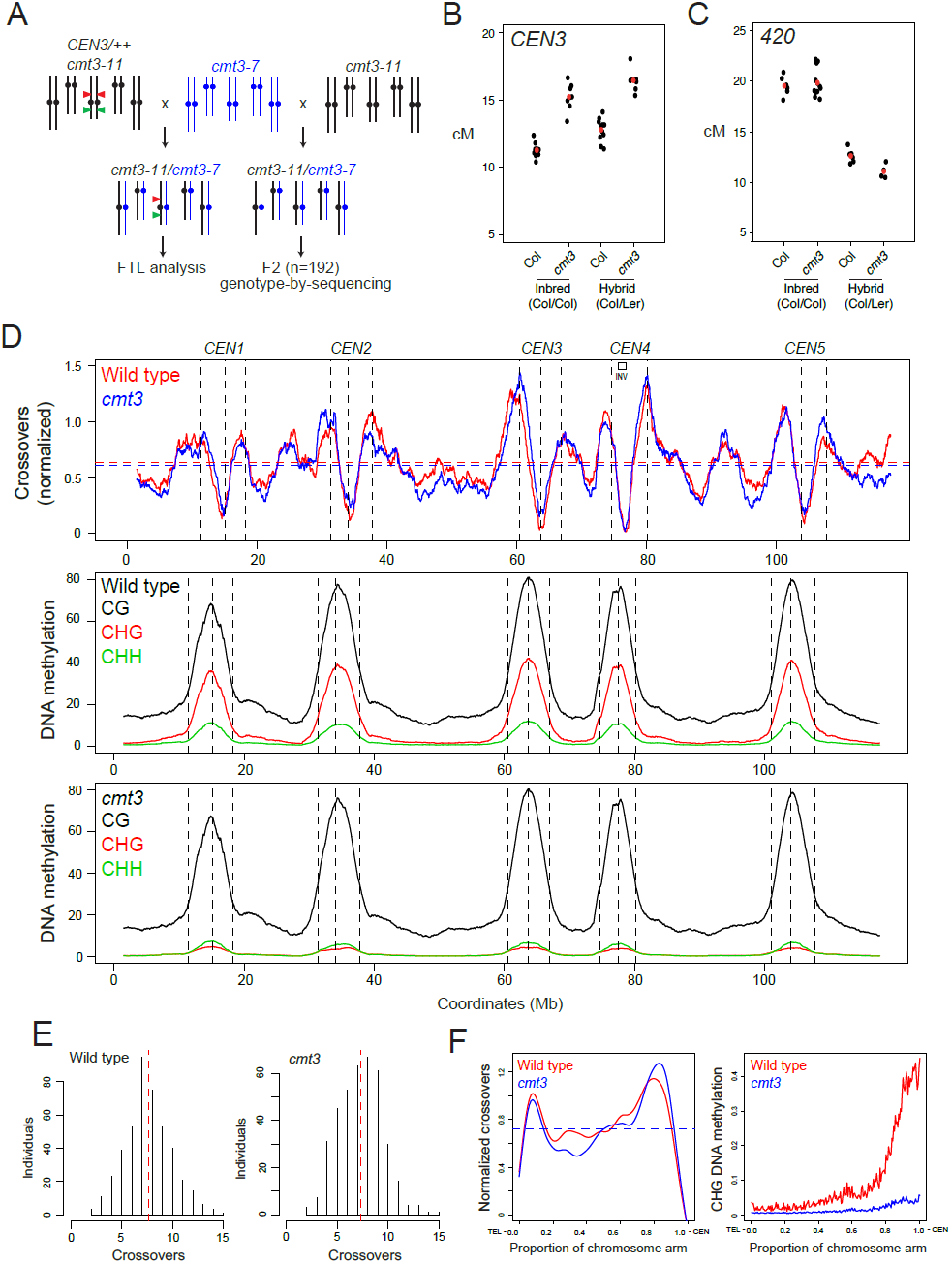
Genome-wide mapping of crossover frequency in *cmt3* non-CG mutants. **(A)** Crossing scheme used to analyse recombination in *cmt3* mutants. Col chromosomes are black and Ler chromosomes are blue. The *CEN3* FTL T-DNAs are indicated by red and green triangles. **(B)** *CEN3* crossover frequency (cM) in wild type and *cmt3*, in Col/Col inbreds, or Col/Ler F_1_ hybrids. Replicate measurements are shown in black and mean values in red. **(C)** Euchromatic *420* crossover frequency in wild type and *cmt3*, in Col/Col inbreds, or Col/Ler F_1_ hybrids, as shown for (B). **(D)** The normalized frequency of crossovers mapped by GBS in wild type (red) and *cmt3* (blue) F_2_ populations is plotted along the Arabidopsis genome, with the 5 chromosomes plotted on a continuous x-axis. Mean values are indicated by horizontal dotted lines. Vertical dotted lines indicate the position of the centromere assembly gaps and the flanking pericentromeric boundaries. Beneath are plots analysing the frequency of CG (black), CHG (red) and CHH (green) context DNA methylation in wild type or *cmt3*, using published BS-seq data (Stroud et al. 2013). The heterochromatic knob inversion on the short arm of chromosome 4 is indicated by the black box and ‘INV’. **(E)** Distribution of crossovers per F_2_ individual for wild type and *cmt3* populations. Red dotted lines indicate mean values. **(F)** Normalized crossover frequency analysed along chromosome telomere (TEL) to centromere (CEN) axes in wild type (red) and *cmt3* (blue) populations. CHG DNA methylation was analysed and plotted and in the same way for wild type (red) or *cmt3* (blue) (Stroud et al. 2013).

### Genome-wide mapping of crossovers in *chromomethylase3* mutants

We next sought to map crossovers genome-wide in wild type compared with a H3K9me2/non-CG mutant background, using segregation of single nucleotide polymorphisms (SNPs). CMT3 is the major CHG context DNA methyltransferase in Arabidopsis, for which mutant alleles are available in both Col (*cmt3-11*) and Ler (*cmt3-7*) backgrounds (Bartee et al. 2001; Lindroth et al. 2001; Stroud et al. 2013, 2014). We therefore generated wild type (Col × Ler) and *cmt3* (*cmt3-11* × *cmt3-7*) F_2_ populations of greater than 700 individuals each (Fig. 3A). To assess centromeric recombination levels in these populations we genotyped Col/Ler simple sequence length polymorphism (SSLP) markers on chromosomes 1 and 3 (Supplemental Table S7). This confirmed significant increases in pericentromeric recombination in the *cmt3* population compared to wild type, consistent with our previous FTL measurements (Supplemental Table S7). To map crossovers at high resolution we performed genotyping-by-sequencing (GBS) of 437 wild type and 384 *cmt3* F_2_ individuals, which identified 3,320 and 2,803 crossovers, respectively (Fig. 3D-3F and Supplemental Table S8) (Choi et al. 2016; Rowan et al. 2015). The crossovers were mapped between Col/Ler SNPs to a mean resolution of 887 bp. The total number of crossovers per wild type F_2_ individual (mean=7.8) was comparable to that observed in similar F_2_ populations (Giraut et al. 2011; Salomé et al. 2012), and was not significantly different in *cmt3* (mean=7.3) (Mann-Whitney-Wilcoxon test, *P*=0.10) (Fig. 3E and Supplemental Table S8).

To analyse crossover distributions throughout the genome we defined, (i) the centromere as the contiguous recombination-silent regions in wild type that surround the centromeric assembly gaps, (ii) the pericentromeres as regions flanking the centromeres with higher than average DNA methylation, and (iii) the chromosome arms as the remainder of the sequence (Fig. 3B and Supplemental Table S9). Consistent with our FTL analysis, we observed that GBS-mapped crossovers were significantly increased in the *cmt3* pericentromeric regions (24.6 versus 27.8% of events were pericentromeric in wild type versus *cmt3, X*^*2*^ *P=*5.60 × 10^−3^), which are strongly depleted of CHG DNA methylation in *cmt3* (Fig 3D, 3F and Supplemental Table S10) (Stroud et al. 2013). We also observed elevated centromeric crossovers (*n*=13) in *cmt3* (*X*^*2*^ *P=*2.63 × 10^−4^), which were completely absent in wild type (Supplemental Table S10). The chromosome arms showed a significant decrease of crossovers in *cmt3* (*X*^*2*^ *P=*1.50 × 10^−3^) (Supplemental Table S10). These data confirm that crossovers increase in proximity to *cmt3* centromeres, but that the increase is strongest in the flanking pericentromeric regions (Fig. 3D and 3F). It is important to note that this experiment is performed in a Col/Ler hybrid context, in contrast to our previous FTL experiments, which are in a Col/Col inbred background (apart from the hemizygous fluorescent protein-encoding transgenes) (Fig. 2). In a Col/Ler hybrid context, centromeric crossovers are likely to be additionally suppressed by structural polymorphisms, including in retrotransposon insertions and *CEN180* repeat copy number and arrangement (Ito et al. 2007; Stuart et al. 2016; Quadrana et al. 2016). For example, the inhibitory effect of structural polymorphism is evident within the ~ 1.17 Mb heterochromatic knob inversion on chromosome 4, which is strongly suppressed for crossovers in both wild type and *cmt3* populations (Fig. 3D) (Fransz et al. 2000, 2016). From these data we conclude that loss of the CHG DNA methyltransferase CMT3 increases meiotic crossover frequency most strongly in the pericentromeric heterochromatin, although structural genetic variation also likely plays an important role in shaping recombination in hybrid contexts.

### Meiotic immunocytology of chromatin and recombination in non-CG/H3K9me2 mutants

To assess H3K9me2 patterns during meiosis we performed immunocytological staining using antibodies against this histone modification and the chromosome axis protein ASYNAPTIC1 (ASY1) (Fig. 4A) (Armstrong et al. 2002). During meiosis *Arabidopsis* centromeres and pericentromeres undergo progressive clustering during prophase-I (Fig. 4A) (Armstrong et al. 2001). We observed that the heterochromatic clusters, detected by DAPI staining of DNA, accumulate dense H3K9me2 throughout leptotene, zygotene and pachytene (Fig. 4A), which are key stages during which meiotic DSB formation and crossover maturation occur (Armstrong et al. 2001; Sanchez-Moran et al. 2007). At leptotene stage when meiotic DSBs occur, H3K9me2 was only detected at background levels in *kyp suvh5 suvh6*, and was reduced in *cmt3* compared to wild type controls (Fig. 4B, 4D and Supplemental Table S11). Similar intermediate reductions in H3K9me2 are observed in *cmt3* somatic cells (Supplemental Fig. S2 and Supplemental Table S12), as previously reported (Yelagandula et al. 2014). Thus H3K9me2 accumulates strongly in *Arabidopsis* heterochromatin during meiosis, and its loss or reduction results in increased pericentromeric crossover.

**Figure 4.**
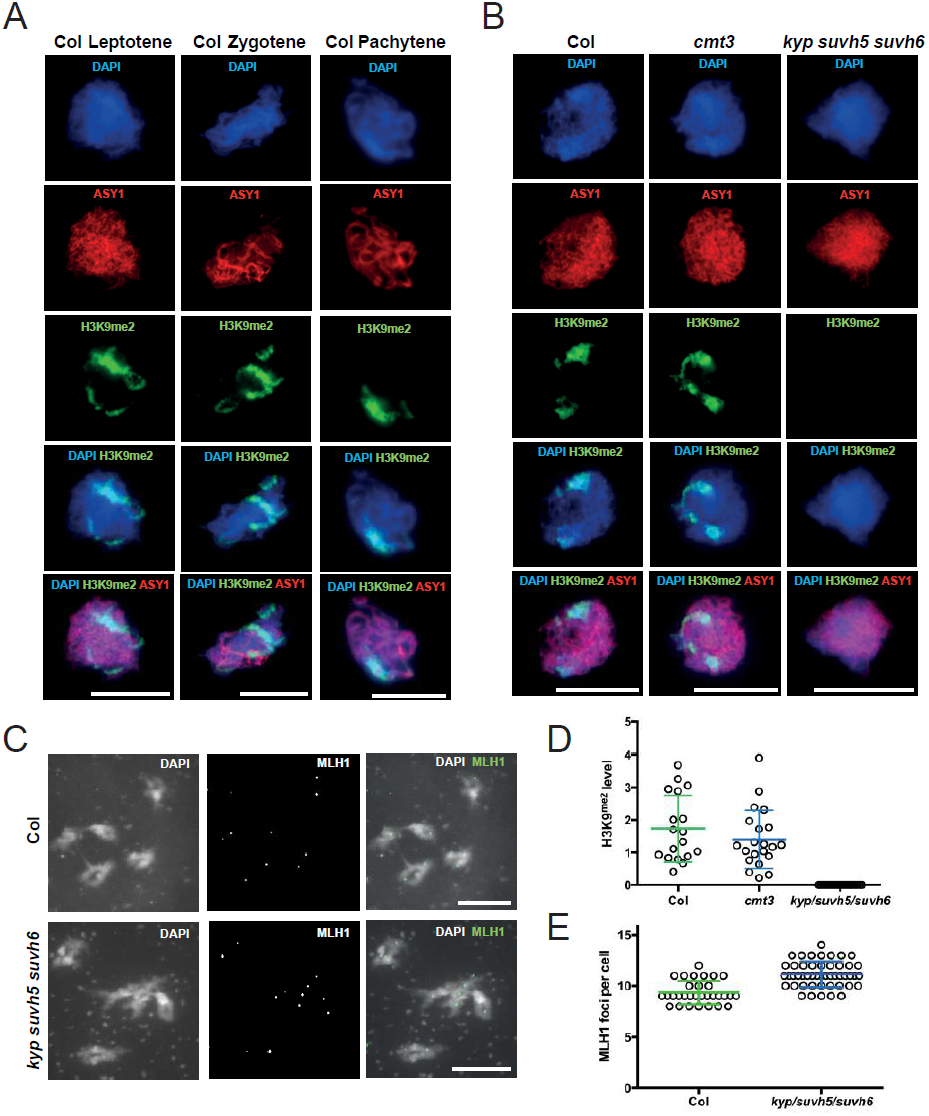
Meiotic heterochromatin is enriched for H3K9me2 and shows increased MLH1 foci in *kyp suvh5 suvh6*. **(A)** Wild type (Col) male meiocytes immunostained for ASY1 (red) and H3K9me2 (green) and DNA (DAPI, blue) at leptotene, zygotene and pachytene stages. The scale bar shows 10 μM. **(B)** As for (A), but showing leptotene stage cells from wild type (Col), *cmt3-11* and *kyp suvh5 suvh6*. (**C)** Male meiocytes at diakinesis stained for DAPI and immunostained for MLH1 (green) in wild type and *kyp suvh5 suvh6*. **(D)** Quantification of H3K9me2 immunostaining in wild type (Col), *cmt3* and *kyp suvh5 suvh6* somatic cells. **(E)** Quantification of MLH1 foci in wild type and *kyp suvh5 suvh6*.

Crossovers can be detected by immunostaining for MLH1, which marks class-I interfering crossover foci (Crismani et al. 2012). Therefore, we scored MLH1 foci associated with euchromatin or heterochromatin, based on DAPI staining, in wild type and *kyp suvh5 suvh6* mutants (Fig. 4C, 4E and Supplemental Table S13). We observed a slight but significant increase in MLH1 foci numbers in *kyp suvh5 suvh6* (mean=11.1) compared to wild type (mean=9.4) at diakinesis (Mann-Whitney-Wilcoxon test, *P* = 1.64 × 10^−7^) (Fig. 4C and 4E and Supplemental Table S13). Importantly, MLH1 foci were also significantly increased in *kyp suvh5 suvh6* heterochromatin (mean=2.9), compared to wild type (mean=1.7) (Mann-Whitney-Wilcoxon test, *P* = 4.22 × 10^−5^) (Fig. 4C and 4E and Supplemental Table S14). This provides cytological support for our crossover mapping and indicates that MLH1-dependent repair contributes to the increase in pericentromeric crossovers observed in H3K9me2/non-CG DNA methylation mutants.

As MLH1 foci were increased in *kyp suvh5 suvh6*, we further investigated the relationship of class I and class II crossover pathways to the observed recombination changes. Approximately 85% of *Arabidopsis* crossovers are dependent on the class I repair pathway, which are interference-sensitive (Mercier et al. 2005; Higgins et al. 2004). Mutants in the class I pathway, for example *zip4*, cause a strong reduction in crossovers and fertility (Chelysheva et al. 2007; Mercier et al. 2005). The *fancm* mutation restores fertility in *fancm zip4* double mutants, by increasing class II non-interfering crossovers (Crismani et al. 2012). Therefore, we constructed *cmt3 zip4* and *cmt3 fancm* double mutants and compared *CEN3* crossover frequency and fertility to wild type and single mutants (Fig. 5A-5C and Supplemental Tables S14 and S15). Unlike *fancm, cmt3* was unable to suppress *zip4* infertility (Fig. 5A-5B and Supplemental Table S15). However, a small but significant *CEN3* crossover increase was observed in *cmt3 zip4*, compared with *zip4* alone (*X*^*2*^ *P=*2.99 × 10^−9^), which indicates that the class II pathway also contributes to increased crossovers in *cmt3* mutant centromeres (Fig. 5C and Supplemental Table S14). Additionally, the *cmt3 fancm* double mutant shows an additive increase in *CEN3* crossover frequency, compared with *cmt3* (*X*^*2*^ *P=*3.34 × 10^−10^) and *fancm* (*X*^*2*^ *P=*2.66 × 10^−7^) single mutants (Fig. 5C and Supplemental Table S14). Hence, increased pericentromeric recombination in H3K9me2 and non-CG DNA methylation mutants involves contributions from both interfering and non-interfering crossover repair.

**Figure 5.**
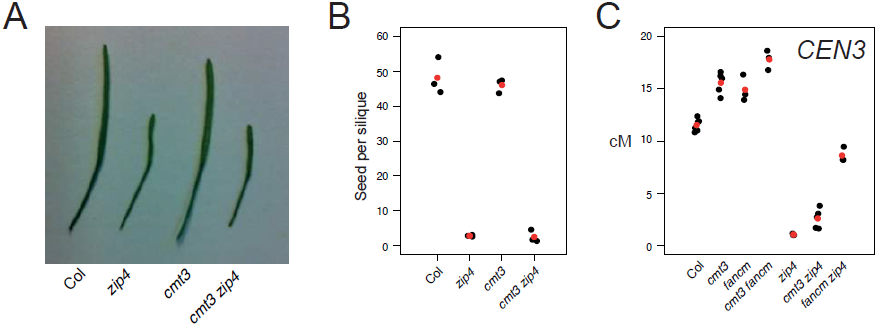
Genetic interactions between class-I and class-II crossover pathways and H3K9me2/non-CG methylation. **(A)** Silique length in wild type (Col), *zip4, cmt3* and *cmt3 zip4*. **(B)** Seed set per silique in wild type (Col), *zip4, cmt3* and *cmt3 zip4.* Replicate measurements are shown in black and mean values in red. **(C)** *CEN3* crossover frequency (cM) in wild type, *cmt3, zip4* and *fancm* mutant backgrounds. Replicate measurements are shown in black and mean values in red.

### Meiotic DSBs are elevated in the centromeres of non-CG/H3K9me2 mutants

During Arabidopsis meiosis, SPO11-1 acts with SPO11-2 and MTOPVI-B to generate DSBs, which can undergo interhomolog repair to form crossovers (Grelon et al. 2001; Hartung et al. 2007; Vrielynck et al. 2016). SPO11 enzymes are related to topoisomerase-VI transesterases and become covalently linked to ~ 20-50 base target site oligonucleotides during DSB formation (Lange et al. 2016; Keeney and Kleckner 1995). We have purified and sequenced Arabidopsis SPO11-1-oligonucleotides in order to map patterns of meiotic DSBs genome-wide, using a complementing *SPO11-1-Myc* line (Choi et al., submitted). In order to profile meiotic DSBs in H3K9me2/non-CG DNA methylation mutants, we crossed *SPO11-1-Myc* to *kyp suvh5 suvh6* and generated SPO11-1-oligonucleotide sequencing libraries (Supplemental Table S16). In wild type, SPO11-1-oligonucleotides are suppressed in centromeric heterochromatin, but are significantly increased in *kyp suvh5 suvh6* mutants (Poisson regression model *P*=1.24 × 10^−14^) (Fig. 6A). Interestingly, the increased DSBs observed in *kyp suvh5 suvh6* were primarily in the centromeres, while crossover increases observed in *cmt3* were primarily in the pericentromeres (Fig. 6A). Hence, while both meiotic DSBs and crossovers increase in H3K9me2/non-CG mutant heterochromatin, they are elevated in adjacent centromeric and pericentromeric regions, respectively.

**Figure 6.**
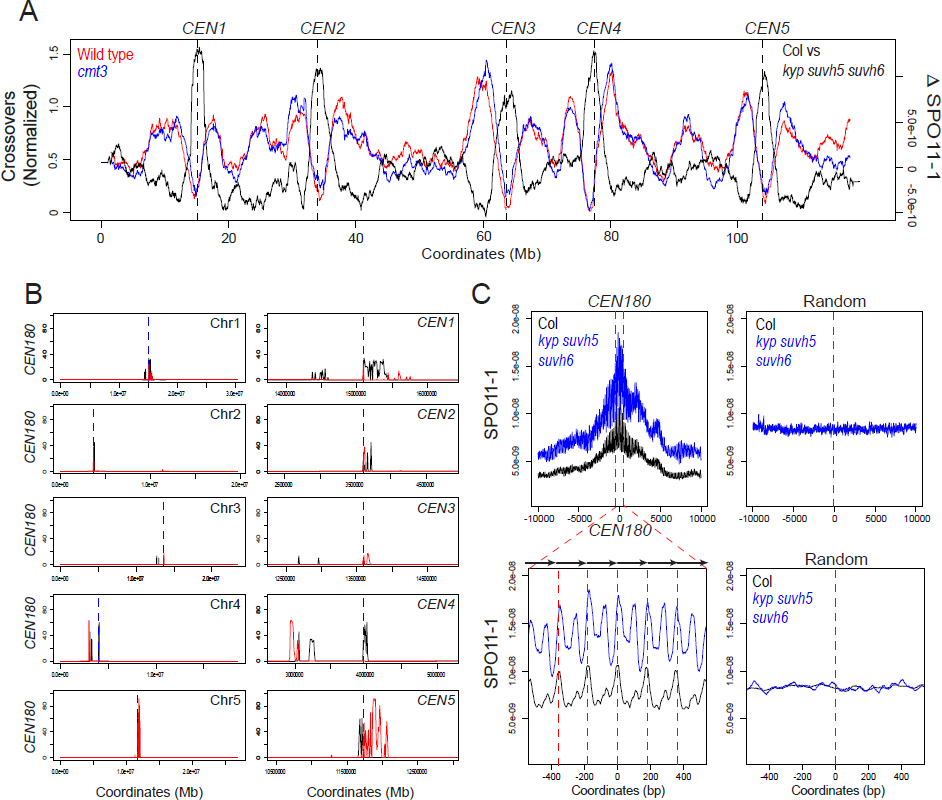
Elevated SPO11-1-oligonucleotides levels in centromeres of *kyp suvh5 suvh6* H3K9me2 mutants. **(A)** The Arabidopsis genome is plotted on a continuous x-axis, showing normalized crossover frequency in wild type (red) and *cmt3* (blue) (as in Fig. 3D), compared with the SPO11-1 differential between wild type and *kyp suvh5 suvh6* (black). The centromere positions are indicated by vertical dotted lines. **(B)** Plots of the Arabidopsis chromosomes showing the density of *CEN180* repeats on forward (black) and reverse (red) strands. The TAIR10 centromeric assembly gaps are shown by the dotted blue line. The plots on the left show whole chromosomes, whereas the plots on the right show a close-up of the centromere. **(C)** The normalized density of SPO11-1-oligos was analysed in 20 kb windows surrounding matches to the *CEN180* consensus in wild type (black) or *kyp suvh5 suvh6* (blue). The lower plot corresponds to a blow-up of a 1 kb windows around the center of the upper plot. The lower plot highlights the approximate positions of *CEN180* repeats using arrows and red dotted lines. Also shown is an identical analysis performed for the same number of randomly chosen sites.

We next analysed DSB frequency around copies of the *CEN180* satellite repeat, which are found in the proximity to the centromeres. Each Arabidopsis chromosome sequence contains a centromere gap, which contains large megabase arrays of *CEN180* repeats (Copenhaver et al. 1999; Kumekawa et al. 2000; Ito et al. 2007; Maheshwari et al. 2017). Further matches to the *CEN180* consensus flank these gaps and we identified 3,397 repeats in the Col reference genome that occur in large tandemly repeated arrays (Fig. 6B). SPO11-1-oligo density was analysed in 20 kb windows around these repeat positions and compared to the same number of randomly chosen windows, in wild type and *kyp suvh5 suvh6* (Fig. 6C). The *CEN180* regions analysed overall showed lower than random SPO11-1-oligonucleotide levels in wild type (Fig. 6C). Consistent with the SPO11-1-oligo increase in centromeric regions at the chromosome scale (Fig. 6A), we observed a pronounced increase in DSBs in the *CEN180* repeats at the fine-scale in *kyp suvh5 suvh6* (Fig. 6C). Interestingly, the SPO11-1-oligo level was highest at the junctions between *CEN180* repeats, though an additional peak internal to the repeats was also observed in *kyp suvh5 suvh6* (Fig. 6C). Together this shows that meiotic DSBs increase in the *CEN180* repeats in H3K9me2 mutant backgrounds. A relatively small number (141) of transposons are transcriptionally upregulated in *kyp suvh5 suvh6* mutants and associated with decreased non-CG methylation (Stroud et al. 2012, 2013). Some of these elements were associated with elevated SPO11-1 oligonucleotides in *kyp suvh5 suvh6* mutants, but others were not (Supplemental Fig. 3), indicating that transcriptional activation is not strictly coupled to upregulation of meiotic DSB formation at Arabidopsis transposons.

## Discussion

Meiotic crossover in proximity to centromeres has been associated with chromosome mis-segregation and aneuploidy in fungi and animals (Rockmill et al. 2006; Lamb et al. 1996; Koehler et al. 1996). Hence, suppression of crossovers within centromeres is thought to play an important role in maintaining fidelity of genome transmission during meiosis. However, abundant evidence for recombination-associated polymorphism in centromeric repeats exists, consistent with the effects of replication slippage, unequal crossover and gene conversion (Wolfgruber et al. 2016; Ma and Bennetzen 2006; Nambiar and Smith 2016). For example, maize centromeric *CRM* retrotransposons have been observed to undergo meiotic gene conversion, but not crossover (Shi et al. 2010), which indicates initiation of meiotic DSBs but with downstream inhibition of recombination steps leading to crossover. We also observe evidence for SPO11-1-dependent DSBs within the Arabidopsis *CEN180* repeats, meaning that meiotic recombination may contribute to polymorphism within centromeric satellite repeat arrays in this species (Ito et al. 2007; Maheshwari et al. 2017).

As centromeric DSBs increase in *kyp suvh5 suvh6* mutants, this demonstrates that epigenetic information, including H3K9me2 and non-CG DNA methylation, plays important roles in suppressing initiation of meiotic recombination in these regions. However, our data also reveal complexity in how chromatin shapes meiotic recombination around plant centromeres. First, while the centromeric regions show increased DSBs in *kyp suvh5 suvh6*, in *cmt3* we observed that crossovers were most elevated in adjacent pericentromeric regions. An important distinction between these experiments is that SPO11-1-oligonucleotides were mapped in a Col/Col homozygous background, whereas mapping crossover necessitates use of polymorphic Col/Ler hybrids. Following DSB formation, resection occurs to generate 3’-single stranded DNA that can perform strand invasion of homologous chromosomes (Keeney and Neale 2006). Interhomolog recombination downstream of strand invasion is sensitive to heterology between the recombining chromosomes, which can have a locally inhibitory effect on crossover formation, and instead promote non-crossover repair and gene conversion (Borts and Haber 1987; Dooner 1986). Arabidopsis centromeric regions exhibit extensive structural variation, including within Gypsy retroelements (Kumekawa et al. 2000; Ito et al. 2007; Stuart et al. 2016; Quadrana et al. 2016), which may therefore suppress centromeric crossover formation downstream of inter-homolog strand invasion, despite activation of meiotic DSBs in non-CG/H3K9me2 mutants. It is also possible that additional chromatin or epigenetic features enriched within the centromeres suppress crossover repair. For example the kinetochore, CENH3 nucleosomes, or further heterochromatic marks such as H2A.W may be differentially enriched within the centromeric versus pericentromeric regions and cause inhibition of crossover maturation (Vincenten et al. 2015; Yelagandula et al. 2014).

An important question raised by our study is why pericentromeric crossover frequency increases in the H3K9me2/non-CG pathway mutants reported here, but not in *met1* and *ddm1* where reduced pericentromeric crossover frequency is observed (Yelina et al. 2015, 2012; Melamed-Bessudo and Levy 2012; Colomé-Tatché et al. 2012; Mirouze et al. 2012). MET1 and DDM1 have major roles in the maintainence of DNA methylation in the CG context. However, their molecular roles in other respects are distinct; (i) non-CG DNA methylation is reduced in *ddm1* to a greater extent than *met1* (Stroud et al. 2013), (ii) gene body methylation is eliminated in *met1* but not *ddm1* (Stroud et al. 2013) and (iii) H3K9me2 is reduced more strongly in *ddm1* compared with *met1* (Deleris et al. 2012; Gendrel et al. 2002). Therefore, we postulate that their common feature, loss of CG methylation within heterochromatin, alters progression and maturation of the meiotic recombination pathway, such that crossovers are favoured in the chromosome arms, at the expense of the pericentromeres. In contrast, H3K9me2/non-CG pathway mutants activate recombination in the heterochromatic regions such that maturation of crossovers in the pericentromeres is increased. Indeed, distinctions in recombination phenotype are consistent with the different effects on transcription, transposition and chromosomal conformation associated with loss of CG versus non-CG DNA methylation maintenance pathways (Stroud et al. 2012; Yelina et al. 2015; Mirouze et al. 2012; Melamed-Bessudo and Levy 2012; Colomé-Tatché et al. 2012; Kato et al. 2003; Miura et al. 2001; Stroud et al. 2013, 2014; Zhang et al. 2006; Lippman et al. 2004; Lister et al. 2008; Henderson and Jacobsen 2008; Feng et al. 2014; Singer et al. 2001).

We propose that while both CG and non-CG DNA methylation inhibit centromeric meiotic DSBs (Choi et al., submitted), only non-CG methylation and/or H3K9me2 inhibit crossovers. In agreement with this idea, euchromatic crossover hotspots in Arabidopsis can be silenced by RNA-directed DNA methylation associated with gain of H3K9me2 and both CG and non-CG methylation (Yelina et al. 2015). Furthermore, in fission yeast arrested recombination intermediates accumulate strongly in wild type heterochromatin, but not in *clr4* and *rik1* (Recombination In K) heterochromatin, which lose H3K9me2 and undergo mitotic and meiotic recombination (Ellermeier et al. 2010; Zaratiegui et al. 2011). Mouse *dnmt3l* mutants also have altered DNA methylation and chromatin signatures and increased DSB initiation within retrotransposons, which is associated with meiotic catastrophe and infertility (Zamudio et al. 2015). In contrast, Arabidopsis H3K9me2 and DNA methylation mutants are fully fertile, despite increased recombination initiation in the centromeric regions, suggesting that increased meiotic DSBs in transposons do not *per se* cause infertility in plants. Suppression of heterochromatic recombination is a major barrier to introducing genetic diversity in crop plants like maize and wheat, where the majority of the chromosome is composed of pericentromeric heterochromatin and that yet contains important genetic variation of functional genes (Choulet et al. 2014; Gore et al. 2009). Therefore, an exciting prospect will be to modulate H3K9me2 and non-CG DNA methylation to unlock pericentromeric mcrossovers in crop breeding programmes.

## Methods

### Plant material

*Arabidopsis* plants were grown under long day conditions (16 hours light/8 hours dark, at 150 μmols light intensity) at 20°C. We used the following mutant alleles; *kyp-6* (SALK_041474) (Chan et al. 2006), *cmt3-11* (SALK_148381) (Chan et al. 2006), *cmt3-7* (Lindroth et al. 2001), *kyp suvh5 suvh6* (SALK_041474, GK-263C05, SAIL_1244_F04) (Johnson et al. 2008), *drm1-2 drm2-2* (SALK_031705, SALK_150863) (Chan et al. 2006), *drm1-2 drm2-2 cmt3-11* (SALK_031705, SALK_150863, SALK_148381) (Chan et al. 2006), *cmt2-3* (SALK_012874) (Stroud et al. 2014), *ddm1* (SALK_000590), *zip4-2* (SALK_068052) (Chelysheva et al. 2007) and *fancm-1* (EMS point mutant) (Crismani et al. 2012). The centromeric FTL lines *CTL5.11* and *LTL5.4* were obtained from the Traffic line population (Wu et al. 2015).

### Mapping of *Ds* insertion sites

Generation of the Ds transposant lines was previously reported. *Ds* insertion sites were amplified by TAIL PCR (Springer et al. 1995; Sundaresan et al. 1995).

### Measuring crossovers using fluorescent pollen and seed

Crossover scoring using the pollen FTL line *CEN3* was performed by flow cytometry, as previously reported (Yelina et al. 2013). Crossover scoring using seed FTL lines (*420, CTL5.11, LTL5.4*) was performed by fluorescent imaging, as previously reported (Ziolkowski et al. 2015a). Statistical analysis of fluorescent count data was performed as described (Ziolkowski et al. 2015b; Yelina et al. 2015).

### Genotyping-by-sequencing

Illumina sequencing libraries were constructed in 96 well format, as previously reported (Rowan et al. 2015; Yelina et al. 2015), with the following minor modifications. DNA was extracted from 3 rosette leaves of 5 week old plants and 150 ng of DNA used as input for each library. DNA shearing was carried out for 20 minutes at 37°C with 0.4U of DNA Shearase (Zymo research). Each set of 96 libraries was sequenced on an Illumina NextSeq500 (300-cycle Mid Output run). Sequencing data was analysed to identify crossovers as previously reported, using the TIGER pipeline (Rowan et al. 2015; Yelina et al. 2015). 384 *cmt3-11* × *cmt3-7 F*_*2*_ individuals were sequenced. Wild type crossovers (CO) were mapped by sequencing 245 Col × Ler F_2_ individuals, which were combined with data from 192 F_2_ individuals (Choi et al. 2016).

The coordinates of crossover intervals called by TIGER were used for subsequent analysis. Centromeres were genetically defined as contiguous regions flanking the TAIR10 centromeric assembly gap that show an absence of crossovers in wild type (Salomé et al. 2012; Copenhaver et al. 1999; Giraut et al. 2011). We define the pericentromeric regions as regions flanking the centromeres with higher than chromosome average levels of DNA methylation. The euchromatic arms constitute the remainder of the chromosomes, from the telomeres to the pericentromeres (Supplemental Table S9). Crossovers (midpoints) were counted in these regions and compared to those expected at random according to physical distance and chi-square tests performed (Supplemental Table S10). The distribution of crossovers along the chromosome telomere-centromere axes was analysed as described (Yelina et al. 2015).

### SPO11-1-oligonucleotide sequencing

A detailed protocol and analysis methodology are provided in an accompanying manuscript (Choi et al.). A complementing *SPO11-1-Myc spo11-1* line was crossed with *kyp suvh5 suvh6* triple mutants, and *SPO11-1-Myc spo11-1 kyp suvh5 suvh6* plants identified for analysis. The *CEN180* consensus sequence (5′ - AACCTTCTTCTTGCTTCTCAAAGCTTTCATGGTGTAGCCAAAGTCCATATGAGTCTTTGGCTTTGTGTCTTCTAACAAGGAAACACTACTTAGGCTTTTAAGATCCGGTTGCGGTTTAAGTTCTTATACTCAATCATACACATGACATCAAGTCATATTCGACTCCAAAACACTAACC-3′) was matched to the TAIR10 reference sequence using the R function matchPattern with max.mismatch set to 90. The coverage value of SPO11-1-oligonucleotides was normalized by total library coverage and analysed in 20 kilobase windows around *CEN180* matches, and compared with analysis of the same number of randomly chosen positions.

### Meiotic immunocytology

Chromosome spreads of Arabidopsis pollen mother cells and immunostaining of ASY1 and H3K9me2 were performed using fresh buds, as described (Armstrong et al. 2002). Immunostaining of MLH1 was performed on acetic acid chromosome spreads on fixed buds, as described (Chelysheva et al. 2010). The following antibodies were used: α-ASY1 (rabbit, 1/500 dilution) (Armstrong et al. 2002), α-H3K9me2 (mouse, 1/200 dilution, Abcam, ab1220) and α-MLH1 (rabbit, 1/200 dilution) (Chelysheva et al. 2010). Microscopy was conducted using a DeltaVision Personal DV microscope (Applied precision/GE Healthcare) equipped with a CDD Coolsnap HQ2 camera (Photometrics). Image capture was performed using SoftWoRx software version 5.5 (Applied precision/GE Healthcare). All slides within an experiment (e.g. Col, *cmt3* and *kyp suvh5 suvh6*) were prepared alongside one another and images captured using the same exposure time. The staining pattern of ASY1 and DNA was used to identify cells at leptotene, zygotene or pachytene stage (Sanchez-Moran et al. 2007). Wild type (Col) cells were first analysed and a threshold pixel intensity value identified that removed background signal. This threshold was then applied to all images prior to further processing. Individual cell images were acquired as Z-stacks of 16 optical sections of 0.25 μm each and the maximum intensity projection of the cell was reconstructed using ImageJ, as described (Lambing et al. 2015). The boundaries of each cell were manually defined and the total signal intensity within the cell measured. An adjacent region outside of the cell was used to measure mean background intensity and this value used to subtract from the within cell intensity. The same methods were used for analysis of meiotic and somatic cells.

### Data Access

SPO11-1-oligonucleotide sequencing data in wild type and *kyp suvh4 suvh5 suvh6* are available at ArrayExpress accession E-MTAB-5041 Username: Reviewer_E-MTAB-5041 Password: MKE8bvew. GBS data from wild type and *cmt3* populations are available at ArrayExpress E-MTAB-5476, Reviewer_E-MTAB-5476, password wv9insP4.

## Acknowledgements

We thank Gregory Copenhaver, Avi Levy and Scott Poethig for FTLs, Raphaël Mercier for *zip4-2 fancm-1*, Steven Jacobsen for *cmt3-7*, Chris Franklin for α-ASY1, Mathilde Grelon for α-MLH1 and the Gurdon Institute for access to microscopes. This work was supported by a William Miller Fellowship from the Watson School of Biological Sciences (CJU), the Howard Hughes Medical Institute-Gordon and Betty Moore Foundation (RAM), as well as grants from the Gatsby Charitable Foundation (IRH), the Royal Society (IRH), BBSRC Meiogenix Industrial Partnership Award (IRH) and by grants from the National Institutes of Health (R01GM067014) and from the National Science Foundation to RAM.

## Author contributions

C.J.U., I.R.H. and R.A.M. designed the study. C.J.U performed all experiments apart from SPO11-1 experiments (K.C.), cytological analysis (C.L) and construction of 171 of the wild type genotyping-by-sequencing libraries (H.S.). Data analysis was performed by C.J.U., C.L., K.C., X.Z., H.S., F.B., J.S., E.E., Y.J., I.R.H. and R.A.M. The manuscript was prepared by C.J.U., I.R.H. and R.A.M.

